# Modelling the response of the PIF plantain seedlings to *Tithonia diversifolia* and clam shells treatments in the nursery

**DOI:** 10.1101/2020.05.22.110320

**Authors:** Cécile Annie Ewané, Thaddée Boudjeko

## Abstract

The seeds availability is the main constraint for the agricultural explosion in sub-Saharan Africa countries. In the case of plantain, there is a lack of seedlings in quantity, but also in quality. The advent of the PIF method was an excellent opportunity to improve the availability of plantain seeds, although the quality is not fully guaranteed. Indeed, the PIF plants produced have posed many problems during the acclimation period indicating a need for solutions to improve their quality. Recent researches done with five treatments using *Tithonia diversifolia* and clam shells have highlighted the improvement of the PIF seedlings quality in terms of growth promotion (biofertilizer action) and protection against black Sigatoka disease (biofungicide action). It seemed essential to determine the best model for robust PIF seedlings. The aim of this study was to analyse the different models that have enabled the production of improved PIF seedlings and to determine the best one. We have modelized the response of PIF seedlings to the different treatment’s protocols. It turns out that the best treatment to apply is T5 (*T. diversifolia* liquid extract), followed by T4 (*T. diversifolia* mulch). However, depending on the expected response in the PIF seedlings, all these treatments have proven to be impactful. *Tithonia diversifolia* liquid extract model is the best and in combination with clams, could be useful to boost the production at low cost and without chemical inputs of large amount of improved vigorous (clean and less susceptible) planting material, impacting thus the food security and poverty alleviation.

## 1. Introduction

Plantain is a staple food that plays a vital role in contributing to food security in Central and West Africa, as well as income generation for millions of people in these regions. Cameroon is ranked 3^rd^ in the world (3.94 millions of tons per year) in terms of plantain production and the first in the Central African Economic and Monetary Community (CEMAC) zone [1], where its consumption is very high. The per capita consumption of plantain result in demand largely outstrips supply provoking very high prices for this commodity on rural, urban and trans-border markets. To meet up with this demand, we need to create new plantations in other to improve the performance of this crop whereas, the creation of these new plantations is difficult because of the problem of unavailability of seedlings in quantity, but also seedlings of quality [2].

Vitroplants are considered as the best and safe seedlings but are not affordable for small poor farmers in sub-Saharan Africa. Thus, farmers are used to plant one sucker to obtain one banana plant as a traditional way of creating banana plants in their plantation and this practice is usually subjected to many diseases and pests. Moreover, the bananas field regeneration is a very slow process with low productivity of viable suckers. An alternative is the ‘plantlet from stem bits’ (PIF), a horticultural propagation method that allows massive production of banana seedlings in just two to three months, in a sanitized environment.

The advent and popularization of the PIF in the 2000s raised hopes for solving the seedlings availability problem [3]. However, after about ten years, the PIF has shown some problems limiting its adoption and are now rejected by some farmers. Indeed, many problems are responsible for plants mortality of about 60% during the establishment of new plantations such as contamination on farmlands and the position of the shoot on explants which influences the vigor of the generated plant [2][4], pest and disease pressure (BSD, banana nematodes and weevil) and declining soil fertility [5]. Indeed, the only control method for BSD in the nursery is leaf removal (deleafing) that seems to be ineffective as seedlings are transplanted to field with 2-4 leaves with high level of black Sigatoka infections, much lower than the recommended 5-6 leaves [6]. The poor smallholder farmers could not buy chemical inputs that are harmful to human and the environment, to improve the performance of the PIF seedlings in nursery and on the farm.

Recent researches have shown that soil amendment with *Tithonia diversifolia* alone or combine to clam shells, *Tithonia diversifolia* mulch, *Tithonia diversifolia* vertical layer and *Tithonia diversifolia* liquid extract improve the growth promotion of the PIF seedlings, and also protects them efficiently against BSD [2]-[4], [7]-[8]. Hence, these treatments seem to act in the improved PIF seedlings production as a vital stimulator (growth promotion and biofungicide actions). There is therefore a need to analyse and classify the best of these different models used in the improvement of PIF seedlings. The aim of this study is to analyse the different models explaining the importance of factors in the production of improved PIF seedlings and to determine the best one. The experiments were conducted in Yaoundé (Cameroon) from September 2014 to March 2017.

## 2. Results

### 2.1. Correlation analysis of the different factors with the PIF seedlings responses to treatments

The variables (treatments and stages) were strongly and significantly correlated (*P*> 0.05) to all the responses (number of shoots, height of shoots, diameter of shoots, area of leaves, BSD severity, total proteins and total polyphenols) of the PIF plantain seedlings. As shown in Table 1, the height and diameter of shoots are positively correlated with treatment T4 and the end stage, and negatively correlated with the initial stage. The BSD severity, area of leaves and number of shoots are negatively correlated with the initial stage and positively correlated with the end stage. The BSD is positively correlated with treatment T3. The total proteins and total polyphenols are both negatively correlated with the treatment T2 and positively correlated with treatment T5, T4 and T5 respectively (Table 1).

**Table 1:** Analysis of correlation between the variables (conditions, treatments and stages) and responses (total proteins, total polyphenols, BSD severity, height of shoots, diameter of shoots, area of leaves and number of shoots). The correlation matrix of Pearson (n) shows positive or negative correlation, but also the strength of the relationship (**bold**). Values in bold are different from 0 with a significance level alpha= 0,05.

| Variables | Number of shoots | Height (cm) | Diameter (mm) | Area of leaves (mm<sup>2</sup>) | BSD Severity (cm<sup>2</sup>) | Total proteins (mg Eq BSA/g FW) | Total polyphenols (mg Eq Cat/g FW) |
| --- | --- | --- | --- | --- | --- | --- | --- |
| Condition-SS | 0,029 | 0,081 | -0,061 | 0,091 | -0,102 | 0,063 | 0,015 |
| Condition-uSS | -0,029 | -0,081 | 0,061 | -0,091 | 0,102 | -0,063 | -0,015 |
| Treatment-T3 | 0,101 | -0,100 | -0,408 | -0,247 | **0,465** | 0,390 | -0,244 |
| Treatment-T1 | -0,264 | -0,499 | -0,384 | 0,348 | -0,286 | -0,044 | -0,273 |
| Treatment-T2 | -0,092 | -0,268 | 0,017 | 0,060 | -0,123 | **-0,634** | **-0,515** |
| Treatment-T4 | 0,242 | **0,597** | **0,577** | -0,068 | 0,182 | -0,250 | **0,535** |
| Treatment-T5 | 0,084 | 0,403 | 0,300 | -0,185 | -0,162 | **0,550** | **0,570** |
| Stage-initial | **-0,871** | **-0,497** | **-0,476** | **-0,692** | **-0,588** | -0,329 | -0,218 |
| Stage-end | **0,871** | **0,497** | **0,476** | **0,692** | **0,588** | 0,329 | 0,218 |

### 2.2. Effect of tested variables on the number of shoots of the PIF plantain seedlings

Regarding the variable tested, type of treatment (T1 to T5), stage of growth (initial at application or end during response evaluation), soil condition (sterile or unsterile), no one had a direct effect on the number of shoots. Concerning combined effects, no models when combined with the sterile condition (Condition-SS) and the unsterile condition (Condition-uSS) significantly affected and positively impacted the number of shoots. The sterile condition and the unsterile condition as well as treatment T4 combined with the duration of the trials (stage-end) significantly and positively impacted the number of shoots (Table 2). Treatments T1 and T2, affected negatively the number of shoot when combined with the duration of production.

**Table 2:**
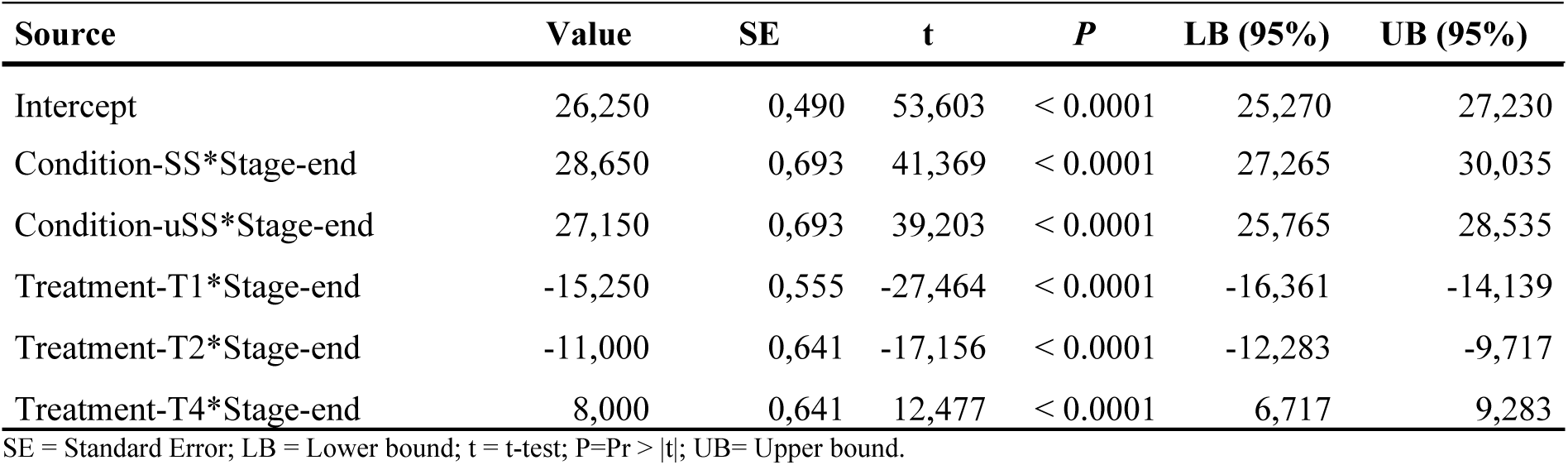
Model parameters for the Number of shoots, obtained from an ANOVA two-ways analysis, showing significant impact of variables (Intercept, conditions, treatments and stages) on the response.

### 2.3. Effect of tested variables on the height of shoots of the PIF plantain seedlings

No variable had a direct effect on the height of shoots. Concerning combined effects, treatments T4 and T5 when combined with the sterile condition significantly and positively affected the height of shoots as well as treatment T4 combined with unsterile condition. On the other hand, treatments T1, T2, T3 and T5 combined with the unsterile condition did not significantly impact the height of shoots. All treatments (T1, T2, T3, T4 and T5) combined with the duration of the trials (stage-end) significantly and positively impacted the height of shoots (Table 3).

**Table 3:**
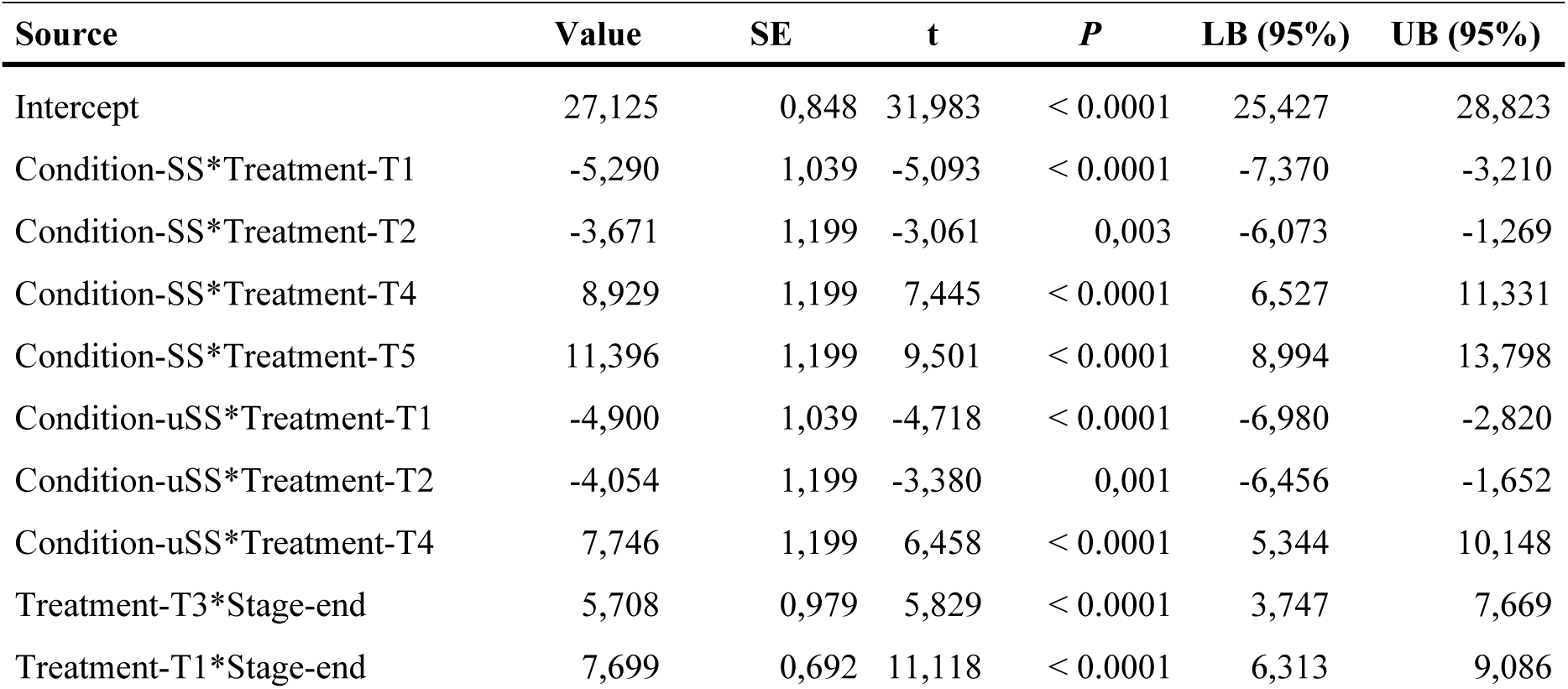
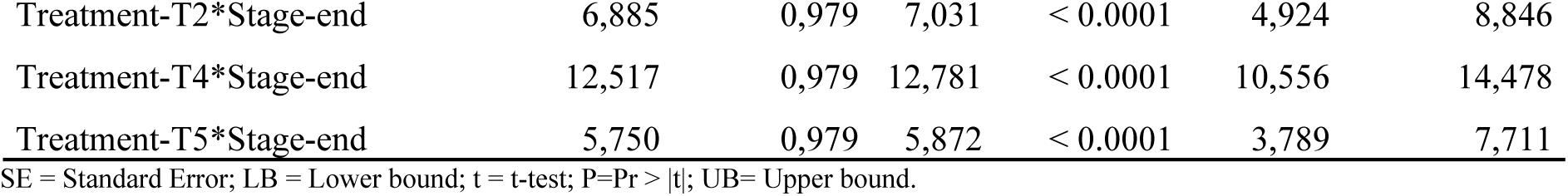
Model parameters for Height of shoots in cm, obtained from an ANOVA two-ways analysis, showing significant impact of variables (Intercept, conditions, treatments and stages) on the response.

| Source | Value | SE | t | P | LB (95%) | UB (95%) |
| --- | --- | --- | --- | --- | --- | --- |
| Intercept | 27,125 | 0,848 | 31,983 | < 0.0001 | 25,427 | 28,823 |
| Condition-SS*Treatment-T1 | -5,290 | 1,039 | -5,093 | < 0.0001 | -7,370 | -3,210 |
| Condition-SS*Treatment-T2 | -3,671 | 1,199 | -3,061 | 0,003 | -6,073 | -1,269 |
| Condition-SS*Treatment-T4 | 8,929 | 1,199 | 7,445 | < 0.0001 | 6,527 | 11,331 |
| Condition-SS*Treatment-T5 | 11,396 | 1,199 | 9,501 | < 0.0001 | 8,994 | 13,798 |
| Condition-uSS*Treatment-T1 | -4,900 | 1,039 | -4,718 | < 0.0001 | -6,980 | -2,820 |
| Condition-uSS*Treatment-T2 | -4,054 | 1,199 | -3,380 | 0,001 | -6,456 | -1,652 |
| Condition-uSS*Treatment-T4 | 7,746 | 1,199 | 6,458 | < 0.0001 | 5,344 | 10,148 |
| Treatment-T3*Stage-end | 5,708 | 0,979 | 5,829 | < 0.0001 | 3,747 | 7,669 |
| Treatment-T1*Stage-end | 7,699 | 0,692 | 11,118 | < 0.0001 | 6,313 | 9,086 |
| Treatment-T2*Stage-end | 6,885 | 0,979 | 7,031 | < 0.0001 | 4,924 | 8,846 |
| Treatment-T4*Stage-end | 12,517 | 0,979 | 12,781 | < 0.0001 | 10,556 | 14,478 |
| Treatment-T5*Stage-end | 5,750 | 0,979 | 5,872 | < 0.0001 | 3,789 | 7,711 |

SE = Standard Error; LB = Lower bound; t = t-test; P=Pr > |t|; UB= Upper bound.

### 2.4. Effect of tested variables on the diameter of shoots of the PIF plantain seedlings

There was no direct effect of the variables observed on the diameter of shoots. Concerning combined effects, treatments T4 and T5 when combined with the sterile condition significantly and positively affected the diameter of shoots, whereas treatments T2, T4 and T5 in unsterile conditions did the same. On the other hand, only treatments T1 combined with the unsterile condition did not significantly impact diameter of shoots. All treatments (T1, T2, T3, T4 and T5) combined with the duration of the trials (stage-end) significantly and positively impacted the diameter of shoots (Table 4).

**Table 4:** Model parameters for Diameter of shoots in mm, obtained from an ANOVA two-ways analysis, showing significant impact of variables (Intercept, conditions, treatments and stages) on the response.

| Source | Value | SE | t | P | LB (95%) | UB (95%) |
| --- | --- | --- | --- | --- | --- | --- |
| Intercept | 2,187 | 0,064 | 33,987 | < 0.0001 | 2,058 | 2,316 |
| Condition-SS*Treatment-T4 | 0,972 | 0,091 | 10,685 | < 0.0001 | 0,790 | 1,154 |
| Condition-SS*Treatment-T5 | 1,208 | 0,091 | 13,277 | < 0.0001 | 1,026 | 1,390 |
| Condition-uSS*Treatment-T3 | -0,189 | 0,074 | -2,551 | 0,013 | -0,338 | -0,041 |
| Condition-uSS*Treatment-T2 | 0,861 | 0,091 | 9,467 | < 0.0001 | 0,679 | 1,044 |
| Condition-uSS*Treatment-T4 | 0,967 | 0,091 | 10,630 | < 0.0001 | 0,785 | 1,149 |
| Condition-uSS*Treatment-T5 | 0,918 | 0,091 | 10,090 | < 0.0001 | 0,736 | 1,100 |
| Treatment-T3*Stage-end | 0,456 | 0,074 | 6,140 | < 0.0001 | 0,307 | 0,605 |
| Treatment-T1*Stage-end | 0,503 | 0,053 | 9,575 | < 0.0001 | 0,398 | 0,608 |
| Treatment-T2*Stage-end | 0,803 | 0,074 | 10,813 | < 0.0001 | 0,655 | 0,952 |
| Treatment-T4*Stage-end | 1,325 | 0,074 | 17,835 | < 0.0001 | 1,176 | 1,474 |
| Treatment-T5*Stage-end | 0,297 | 0,074 | 3,993 | 0,000 | 0,148 | 0,445 |

SE = Standard Error; LB = Lower bound; t = t-test; P=Pr > |t|; UB= Upper bound.

### 2.5. Effect of tested variables on the area of leaves of the PIF plantain seedlings

No variable had a direct effect on the area of leaves. Concerning the combined effects, treatments T1 and T2 when combined with the sterile condition significantly affected the area of leaves. On the other hand, only treatments T3, T4 and T5 combined with the unsterile condition did not significantly impact the area of leaves. To positively impact the area of leaves, there were treatments T1 and T2 in sterile condition and treatments T2, T4 and T5 in the unsterile condition. treatments T1, T2, T3 and T5 combined with the duration of the trials (stage-end) significantly and positively impacted the area of leaves (Table 5).

**Table 5:** Model parameters for Area of leaves in mm^2^, obtained from an ANOVA two-ways analysis, showing significant impact of variables (Intercept, conditions, treatments and stages) on the response.

| Source | Value | SE | t | P | LB (95%) | UB (95%) |
| --- | --- | --- | --- | --- | --- | --- |
| Intercept | 1342,467 | 216,792 | 6,192 | < 0.0001 | 908,349 | 1776,585 |
| Condition-SS*Treatment-T1 | 2558,021 | 265,514 | 9,634 | < 0.0001 | 2026,337 | 3089,704 |
| Condition-SS*Treatment-T2 | 1385,583 | 306,590 | 4,519 | < 0.0001 | 771,648 | 1999,518 |
| Condition-uSS*Treatment-T1 | 2134,296 | 265,514 | 8,038 | < 0.0001 | 1602,612 | 2665,979 |
| Condition-uSS*Treatment-T2 | 1669,716 | 306,590 | 5,446 | < 0.0001 | 1055,781 | 2283,652 |
| Treatment-T3*Stage-end | 513,776 | 250,329 | 2,052 | 0,045 | 12,500 | 1015,052 |
| Treatment-T1*Stage-end | 2987,417 | 177,010 | 16,877 | < 0.0001 | 2632,961 | 3341,872 |
| Treatment-T2*Stage-end | 2193,317 | 250,329 | 8,762 | < 0.0001 | 1692,041 | 2694,593 |
| Treatment-T5*Stage-end | 715,843 | 250,329 | 2,860 | 0,006 | 214,567 | 1217,119 |

SE = Standard Error; LB = Lower bound; t = t-test; P=Pr > |t|; UB= Upper bound.

### 2.6. Effect of tested variables on the BSD severity of the PIF plantain seedlings

BSD severity was not directly impacted by none of the variables studied. Concerning the combined effects, treatments T1, T2 and T5 when combined with the sterile condition significantly affected the BSD severity. On the other hand, treatment T5 combined with the unsterile condition did not significantly impact the BSD severity. To positively impact the BSD severity, there were treatments T1, T2 and T3 in the sterile conditions and treatments T1, T2, T3 and T4 in the unsterile condition. All the treatments (T1, T2, T3, T4 and T5) combined with the duration of the trials (stage-end) significantly and positively impacted the BSD severity (Table 6). Since our target is to negatively impact BSD severity and that non of the combination did it, from table 6, the following group of combination can be seen as having a less favourable impact on BSD severity (treatments T1, T2 and T5 combined to sterile conditions; treatments T1 and T2 combined to unsterile conditions) and treatments T1 and T5 combined with stage-end).

**Table 6:** Model parameters for BSD Severity in cm^2^, obtained from an ANOVA two-ways analysis, showing significant impact of variables (Intercept, conditions, treatments and stages) on the response.

| Source | Value | SE | t | P | LB (95%) | UB (95%) |
| --- | --- | --- | --- | --- | --- | --- |
| Intercept | 30,750 | 6,134 | 5,013 | < 0.0001 | 18,468 | 43,032 |
| Condition-SS*Treatment-T1 | 21,450 | 7,512 | 2,855 | 0,006 | 6,407 | 36,493 |
| Condition-SS*Treatment-T2 | 26,890 | 8,674 | 3,100 | 0,003 | 9,520 | 44,260 |
| Condition-SS*Treatment-T5 | 22,138 | 8,674 | 2,552 | 0,013 | 4,768 | 39,507 |
| Condition-uSS*Treatment-T3 | 48,675 | 7,082 | 6,873 | < 0.0001 | 34,493 | 62,857 |
| Condition-uSS*Treatment-T1 | 27,325 | 7,512 | 3,637 | 0,001 | 12,282 | 42,368 |
| Condition-uSS*Treatment-T2 | 21,885 | 8,674 | 2,523 | 0,014 | 4,515 | 39,255 |
| Condition-uSS*Treatment-T4 | 33,113 | 8,674 | 3,817 | 0,000 | 15,743 | 50,482 |
| Treatment-T3*Stage-end | 205,325 | 7,082 | 28,991 | < 0.0001 | 191,143 | 219,507 |
| Treatment-T1*Stage-end | 23,025 | 5,008 | 4,598 | < 0.0001 | 12,997 | 33,053 |
| Treatment-T2*Stage-end | 39,385 | 7,082 | 5,561 | < 0.0001 | 25,203 | 53,567 |
| Treatment-T4*Stage-end | 125,450 | 7,082 | 17,713 | < 0.0001 | 111,268 | 139,632 |
| Treatment-T5*Stage-end | 28,400 | 7,082 | 4,010 | 0,000 | 14,218 | 42,582 |

SE = Standard Error; LB = Lower bound; t = t-test; P=Pr > |t|; UB= Upper bound.

### 2.7. Effect of tested variables on the total protein contain of the PIF plantain seedlings

No variable had a direct effect on the total proteins. Concerning the combined effects, treatments T1, T2, T4 and T5 when combined with the sterile condition significantly affected the total proteins. On the other hand, treatment T5 combined with the unsterile condition did not significantly impact the total proteins. To significantly and positively impact the total proteins, there were treatments T5 in the sterile condition, treatment T3 on the unsterile condition and treatments T1, T3, T4 and T5 combined with the duration of the trials (stage-end) (Table 7).

**Table 7:** Model parameters for Total Proteins in mg Eq BSA per g of FW, obtained from an ANOVA two-ways analysis, showing significant impact of variables (Intercept, conditions, treatments and stages) on the response.

| Source | Value | SE | t | P | LB (95%) | UB (95%) |
| --- | --- | --- | --- | --- | --- | --- |
| Intercept | 6.227 | 0.367 | 16.987 | < 0.0001 | 5.493 | 6.961 |
| Condition-SS*Treatment-T1 | -2.398 | 0.449 | -5.341 | < 0.0001 | -3.297 | -1.499 |
| Condition-SS*Treatment-T2 | -5.963 | 0.518 | -11.503 | < 0.0001 | -7.002 | -4.925 |
| Condition-SS*Treatment-T4 | -4.036 | 0.518 | -7.786 | < 0.0001 | -5.074 | -2.998 |
| Condition-SS*Treatment-T5 | 3.658 | 0.518 | 7.056 | < 0.0001 | 2.620 | 4.696 |
| Condition-uSS*Treatment-T3 | 0.880 | 0.423 | 2.078 | 0.042 | 0.032 | 1.727 |
| Condition-uSS*Treatment-T1 | -2.246 | 0.449 | -5.003 | < 0.0001 | -3.145 | -1.347 |
| Condition-uSS*Treatment-T2 | -5.980 | 0.518 | -11.535 | < 0.0001 | -7.018 | -4.942 |
| Condition-uSS*Treatment-T4 | -3.582 | 0.518 | -6.909 | < 0.0001 | -4.620 | -2.544 |
| Treatment-T3*Stage-end | 3.269 | 0.423 | 7.722 | < 0.0001 | 2.421 | 4.116 |
| Treatment-T1*Stage-end | 2.347 | 0.299 | 7.840 | < 0.0001 | 1.747 | 2.946 |
| Treatment-T4*Stage-end | 1,886 | 0,423 | 4,455 | < 0.0001 | 1,038 | 2,733 |
| Treatment-T5*Stage-end | 3,527 | 0,423 | 8,332 | < 0.0001 | 2,679 | 4,374 |

SE = Standard Error; LB = Lower bound; t = t-test; P=Pr > |t|; UB= Upper bound.

### 2.8. Effect of tested variables on the total polyphenol contain of the PIF plantain seedlings

Only combined effects were observed. Treatments T2, T4 and T5 when combined with the sterile condition of growth (Condition-SS) significantly affected the total polyphenols. On the other hand, treatment T1 combined with the unsterile condition did not significantly impact the total polyphenols. To positively impact the total polyphenols, there were treatments T4 and T5 in the sterile conditions and on the unsterile condition. Only treatments T4 and T5 combined with the duration of the trials (stage-end) significantly and positively impacted the total polyphenols (Table 8).

**Table 8:** Model parameters for Total Polyphenols in mg Eq Cat per g of FW, obtained from an ANOVA two-ways analysis, showing significant impact of variables (Intercept, conditions, treatments and stages) on the response.

| Source | Value | SE | t | P | LB (95%) | UB (95%) |
| --- | --- | --- | --- | --- | --- | --- |
| Intercept | 3.575 | 0.721 | 4.955 | < 0.0001 | 2.130 | 5.020 |
| Condition-SS*Treatment-T2 | -3.537 | 1.020 | -3.466 | 0.001 | -5.580 | -1.494 |
| Condition-SS*Treatment-T4 | 5.647 | 1.020 | 5.535 | < 0.0001 | 3.604 | 7.690 |
| Condition-SS*Treatment-T5 | 5.214 | 1.020 | 5.110 | < 0.0001 | 3.171 | 7.257 |
| Condition-uSS*Treatment-T3 | -2.237 | 0.833 | -2.685 | 0.009 | -3.905 | -0.568 |
| Condition-uSS*Treatment-T2 | -3.532 | 1.020 | -3.462 | 0.001 | -5.575 | -1.489 |
| Condition-uSS*Treatment-T4 | 5.870 | 1.020 | 5.753 | < 0.0001 | 3.826 | 7.913 |
| Condition-uSS*Treatment-T5 | 5.376 | 1.020 | 5.269 | < 0.0001 | 3.333 | 7.419 |
| Treatment-T4*Stage-end | 3.759 | 0.833 | 4.512 | < 0.0001 | 2.091 | 5.427 |
| Treatment-T5*Stage-end | 5.425 | 0.833 | 6.512 | < 0.0001 | 3.757 | 7.093 |

SE = Standard Error; LB = Lower bound; t = t-test; P=Pr > |t|; UB= Upper bound.

Globally, taking into consideration the positive impacts of the different combined factors on studied responses, it can be observed only treatment T5 combined to the duration of the trial (stage-end) enhanced 6 responses of the 7 measured, followed by treatment T1 combined to duration of trial and sterile condition combined to treatment T5 (5 over 7). Moreover, the factors combinations that less enhanced the BSD severity were sterile and unsterile conditions respectively combined to treatments T1 and T2.

### 2.9. Principal Components Analysis (PCA)

From the PCA two-dimensions, Factor 1 which represented 50.63% of the variability was most influenced by height of shoots, diameter of shoots and number of shoots, while Factor 2, representing 16.78%, was mainly impacted by area of leaves and total polyphenols. BSD severity mostly imparted Factor 3 (16.36%) and in a certain degree F1, F2 and F4. Total polyphenols mostly impacted F5 while total proteins mostly impacted F4 (Table 11). The PCA two-dimensions representation according to F1 and F2 of all the variables and observations, clearly show the different groups and spatial distributions (Figure 1). The group consisted mostly of samples at the end stage who received T1 and T3 treatments in the upper right quarter, with positive F1 and F2 coordinates are influenced by the parameter, area of leaves, number of shoots and BSD severity. On the other hand, the second clear group consisted of samples that received treatments T4 and T5 combined to end stage was located in the down right quarter with positive F1 and negative F2. This group was influenced by parameters diameter of shoots, height of shoots, total protein and total polyphenol.

**Table 9:** Dependent variables weight on the different factors obtained through Principal Component Analysis (PCA).

|  | F1 | F2 | F3 | F4 | F5 |
| --- | --- | --- | --- | --- | --- |
| Total proteins | 6.933 | 4.559 | 33.168 | 37.725 | 12.097 |
| Total polyphenols | 16.254 | 22.193 | 2.536 | 4.327 | 52.142 |
| BSD Severity | 10.379 | 14.151 | 26.322 | 16.925 | 3.197 |
| Height of shoots | 24.270 | 3.422 | 1.972 | 0.761 | 15.304 |
| Diameter of shoots | 18.590 | 0.313 | 21.282 | 4.962 | 13.487 |
| Area of leaves | 1.601 | 44.286 | 13.025 | 34.247 | 0.460 |
| Number of shoots | 21.973 | 11.075 | 1.695 | 1.053 | 3.314 |

**Table 10:** Experimental design for the study of the responses of plantain PIF seedlings for different *Tithonia diversifolia* and clam shells models.

| Completely Randomized Block Device |  |  |
| --- | --- | --- |
|  | Greenhouse | Shade |
| Phase | Germination | Acclimatization |
| Purpose | Production of the PIF seedlings | Survey of the seedling's growth |
| Experimental unit (EU) | Each treatment | Each treatment |
| Substrate to amend | Sawdust | Black soil and sand |
| Number of plants/EU | Three (03) Explants | At least three (3) plants |
| Container | Propagator | Plastic planter bags |
| Block | A sterilized substrate block (B1) | A non-sterilized substrate block (B2) |
| Treatment number | Five (05) in Controlled Condition | Five (05) in Uncontrolled Condition |
| Condition | Sterile Substrate (SS-Industrial) | unSterile Substrate (uSS-Farmer one) |
| Treatment | 1. Clam shells 1% (T1)_[2]<br>2. Clam shells and *T. diversifolia* (T2)_[4]<br>3. One vertical layer *T. diversifolia* flakes (T3)_[7]<br>4. 4 cm Mulch layer of *T. diversifolia* (T4)_[3]<br>5. *T. diversifolia* Liquid extract of 15 days (T5)_[8] |  |
| Variable | Conditions<br>Treatments<br>Stages |  |
| Response | Total proteins<br>Total polyphenols<br>BSD severity<br>Height of shoots<br>Diameter of shoots<br>Area of leaves<br>Number of shoots |  |
| Stage | Initial<br>End |  |

**Table 11:** Presentation of the definition of the initial stage and end stage of the different responses of plantain PIF seedlings and the reference of assessment method.

| Response | Initial Stage | End Stage | Assessment method |
| --- | --- | --- | --- |
| Number of shoots | The day the germination started in the greenhouse | 35 days after the start of germination in the greenhouse | [2] |
| Height of shoots | The day the seedlings were weaned and put in the shade | 42 days after weaning in the shade | [2] |
| Diameter of shoots | The day the seedlings were weaned and put in the shade | 42 days after weaning in the shade | [2] |
| Area of leaves | The day the seedlings were weaned and put in the shade | 42 days after weaning in the shade | [2, 26] |
| BSD severity | The day the leaves were inoculated with *M. fijiensis* | 12 days after the inoculation of leaves with *M. fijiensis* | [2] [27-28] |
| Total proteins | The before inoculation stage | The post-inoculation stage | [29] |
| Total polyphenols | The before inoculation stage | The post-inoculation stage | [30] |

**Figure 1:**
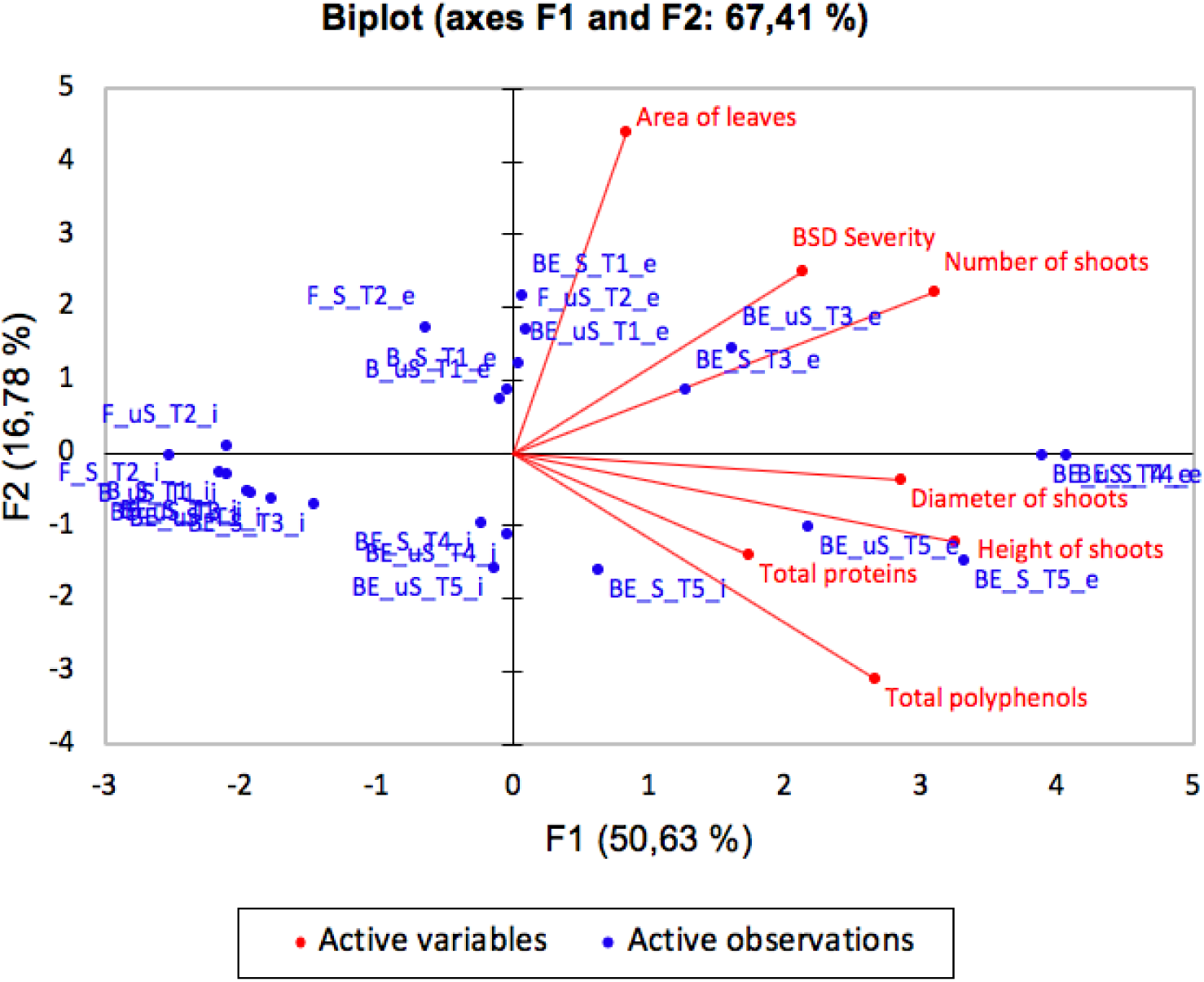
Principal components Analysis (PCA) two-dimensions representation according to F1 and F2 of all the variables and observations, showing different groups and spatial distributions.

Factor 3 have quite the same percentage of explained data variability as factor 2. In this regard, the spatial representation of F1 vs F3 permit to observe different clusters. Hence, the PCA two-dimensions representation according to F1 and F3 of all the variables and observations, clearly show the dissimilarity between the groups and their spatial distributions, but also revealed homogenous groups (Figure 2). The first cluster consisted mostly of samples at the end stage who received T3 and T5 treatments in the upper right quarter, with positive F1 and F2 coordinates are influenced by the parameter, total protein, number of shoots and BSD severity. The second cluster consisted of samples that received treatments T4 and T1 combined to end stage was located in the down right quarter with positive F1 and negative F2. This group was influenced by parameters diameter of shoots, height of shoots, area of leaves and total polyphenols.

**Figure 2:**
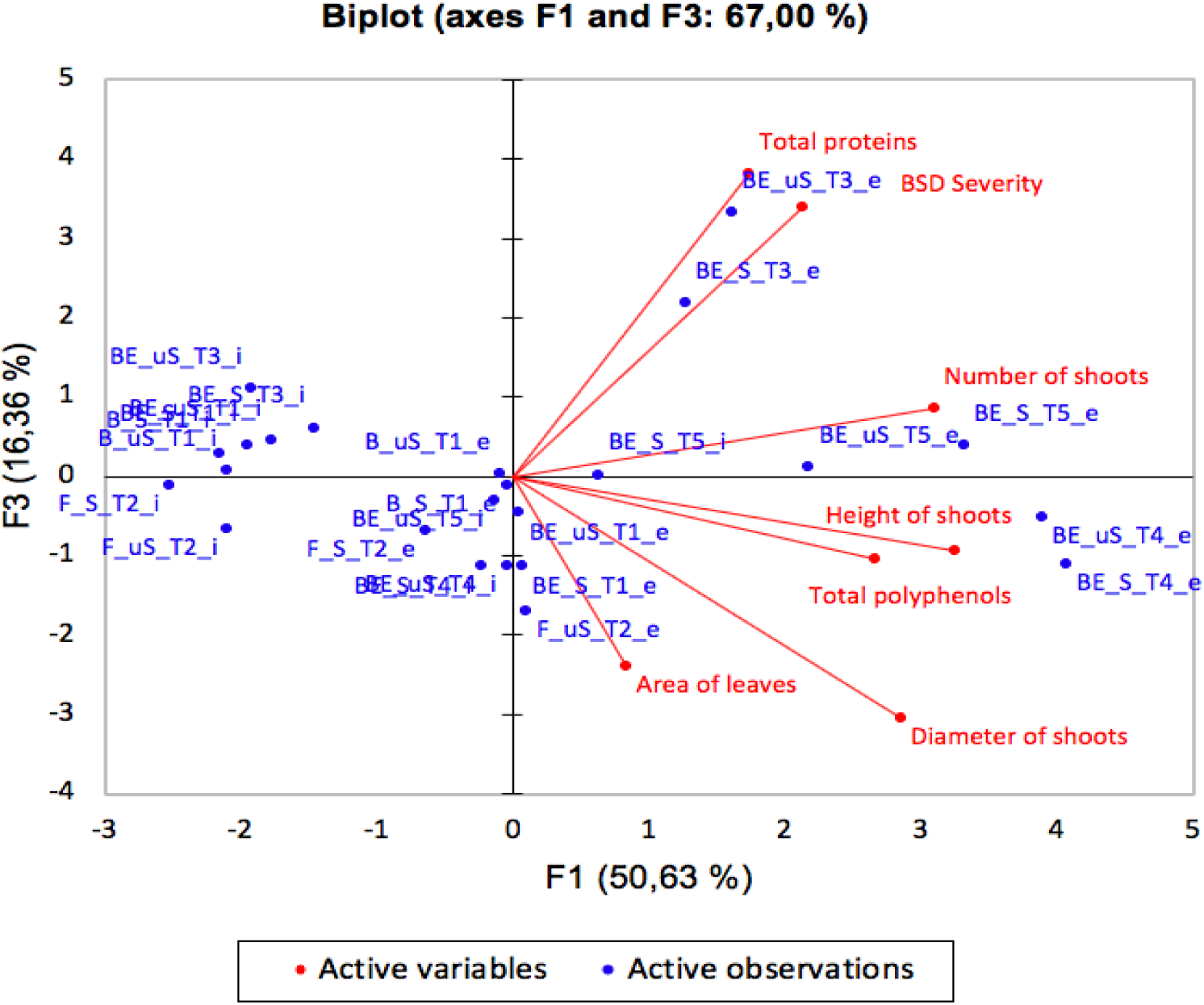
Principal components Analysis (PCA) two-dimensions representation according to F1 and F3 of all the variables and observations, showing different groups and spatial distributions.

## 3. Discussion

The aim of this study was to analyse the different models that have enabled the production of improved PIF seedlings and to determine the best one. Two of these treatments T5 and T4 have been identified as overall impacting mostly the PIF plantain seedlings responses in the greenhouse and the shade. Indeed, the *T. diversifolia* liquid extract (T5) and *T. diversifolia* mulch (T4) have shown growth promotion and antifungal activities in the PIF seedlings [3][8] as well as the other treatments (T3, T4 and T5) despite the less global impact [2][4][7]. The five models based on clam shells and *T. diversifolia* are organic matter that have been shown to activate the growth promotion and natural defense systems of plants through the increase synthesis of nutrients and defensive metabolites [9]-[10]. The organic matter provides nutrients to plants which participate in osmotic regulation, cellular permeability, and may act as structural components and essential metabolites of growth and development [11]; but also, defensive metabolites acting in plant such as the biofungicide effect of organic matter highlighted on the susceptible *Musa* spp. against BSD [12].

Depending on the expected response in the PIF seedlings, the five models are impacting. The increase of the number of shoots is positively impacted by all the models combined with both conditions. Indeed, the abundant shoots’ growth on the suckers is related to the activity of the apical meristem generation favoured by the nitrogen contain in *T. diversifolia* which is involved in division and enlargement of cells in the apical meristem [13]. The height and the diameter of shoots are positively impacted in both conditions by treatments T4 and T5 based on *T. diversifolia*, commonly known acting as plant organic fertilizer in many plants [14]-[16]. Furthermore, *T. diversifolia* tissues are mainly composed of 3-5% nitrogen, 0.5-2.5% phosphorus and 4-6% potassium [17]-[18], mineral elements deeply involved in plant growth promotion. The area of leaves is impacted regardless of the condition by treatments T1 and T2 both containing clam shells. Indeed, clam shells are a rich source of chitin and derivatives that have been shown to influence on growth promoting components, precisely the chitin direct action as fertilizer due to his high nitrogen content and low carbon-nitrogen ratio (C/N) [9]-[10].

The BSD severity is impacted by all the five models, with the most impacting being treatment T2 in the sterile condition and T3 in the unsterile condition. Indeed, *T. diversifolia* is acting as a fungicide in the control of many culture due to the secondary metabolites it contains [19]- [20], while clam shell provides an excellent protection against plant diseases [9]. The total proteins are impacted with treatments T5 and T3 in the sterile condition and the unsterile condition respectively, while the total polyphenols are impacted in both conditions by treatments T4 and T5. These models are based on *T. diversifolia* known as a promoter of natural defensive systems (synthesis of nutrients and defensive metabolites) in plants [9]. Two essential elements in *Tithonia diversifolia* could explant this models’ impact on total proteins and total polyphenols. Nitrogen involved in the preparation of macromolecules and potassium known as an activator of different enzymes [11][21] notably the phenylalanine ammonia lyase (PAL), involved in the biosynthesis of the polyphenol compounds in plants [22]- [23].

Overall, the treatment T5 is the most impacting one for the production of the improved PIF plantain seedlings in the nursery. It is based on *Tithonia diversifolia* liquid extract, and act as a fertilizer and fungicide in the control of disease of PIF seedling as previously reported for another pathosystem [14][20]. However, the impactful action of treatments T1 and T2 on the area of leaves and on the BSD severity in both conditions should be considered in a combined treatment model of *Tithonia diversifolia* liquid extract and clam shells for more improvement of PIF plantain seedlings vigor. Since, the fermented chitin waste (FCW) have been recently shown to enhance the lettuce and rice performance by acting as a plant growth stimulator [24]-[25]. Further studies using this treatment T5 are needed to (1) understand the molecular mechanisms underlying the relationship between the improved PIF seedling and the *Tithonia diversifolia* liquid extract, (2) evaluate this liquid extract effect on other bananas diseases and pests, as well as on other plants, and (3) to position the improved PIF vis-à-vis the vitroplants known as the best banana seeds.

## 4. Materials and Methods

### 4.1. Plant materials and Substrates

Plantain suckers (*Musa* spp., genome AAB) were collected from farms in the centre region of Cameroun. The clam shells were collected from the municipality of Mouanko, while *T. diversifolia* tissues were obtained from farmlands around Yaoundé (Cameroon). The causal agent of black Sigatoka disease (BSD) was provided by the African Centre for Research on Bananas and Plantains (CARBAP-Cameroon). The sawdust, sand and black soil used to formulate the PIF substrates were collected and sterilized at different temperatures and time intervals as previously described by Ewané *et al*. (2019). The PIF substrate in the greenhouse was the sawdust while it was the sand and the black soil (1/3 and 2/3) in the shade.

### 4.2. Experimental Design and Evaluation of different PIF seedlings responses

The experiments design of this study and the method used are presented in Table 10. The variables (conditions, treatments and stages) and responses (number of shoots, height of shoots, diameter of shoots, area of leaves, BSD severity, total proteins and total polyphenols) were evaluated at the initial stage and at the end stage and presented in Table 11. The number of shoots was count, the height of shoots, diameter of shoots, area of leaves and BSD severity measured, total proteins and total polyphenols quantified as described by [2-4] [7-8].

### 4.3. Statistical Analyses

The different treatment responses (number of shoots, height of shoots, diameter of shoots, area of leaves, BSD severity, total proteins and total polyphenols) were analysed by performing a two-way ANOVA with XLSTAT software [31]. Each plant being taken as experimental unit, and stage and treatment as factors. Principal components analysis (PCA) with Pearson correlation between the different variables was also performed with XLSTAT software.

## Acknowledgments

We thank Sylvain SADO for his help in data analysis and stimulating discussions. We also thanks Oscar NGUIDJO for his permanent help and support.

## References

1. FAO. Food and Agriculture Organization of the United Nations. FAO Statistics: Banana. 2018. http://www.fao.org/faostat/en/#data/QC

2. Ewané, C.A.; Ndongo, F.; Ngoula, K.; Tene Tayo; P.M., Opiyo, S.O. and Boudjeko, T. Potential biostimulant effect of clam shells on growth promotion of plantain PIF seedlings (*var*. Big Ebanga & Batard) and relation to black Sigatoka disease susceptibility. American Journal of Plant Science 2019, 10, 1763–1788.

3. Meshuneke, A.; Ewané, C.A.; Tatsegouock R.N.; Boudjeko, T. *Tithonia diversifolia* mulch stimulates the growth of plantain PIF seedlings and induces a less susceptibility to *Mycosphaerella fijiensis* in the nursery. American Journal of Plant Science 2020, 11, 672–692.

4. Ewané, C.A.; Milawé, C.A.; Ndongo, E.F.; Boudjeko, T. Influence of clam shells and Tithonia diversifolia powder on growth of plantain PIF seedlings (var. French) and their sensitivity to Mycosphaerella fijiensis. African Journal of Agricultural research 2020, 15, 393–411.

5. Lefranc, L.M.; Lescot, T.; Staver, C.; Kwa, M.; Michel, I.; Nkapnang, I.; Temple, L. Macroprop-agation as an Innovative Technology: Lessons and Observations from Projects in Cameroon. Acta Horticultural 2010, 8, 727–734.

6. Nfor, T.D.; Ajong, F.D.; Nuincho, L.I. Evaluation of varietal response to black Sigatoka caused by *Mycosphaerella fijiensis* Morelet in banana nursery. International Research Journal of Plant Science 2011, 2, 299–304.

7. Ewané, C.A.; Meshuneke, A.; Tatsegouock R.N.; Boudjeko T. Vertical layers of *Tithonia diversifolia* flakes amendment improves plantain seedling performance. American Journal of Agricultural Research. 2020, 5. https://10.28933/ajar-2020-03-2905

8. Tatsegouock, R.N.; Ewané, C.A.; Meshuneke, A.; Boudjeko, T. Plantain bananas PIF seedlings treatment with liquid extracts of *Tithonia diversifolia* induces resistance to black Sigatoka disease. American Journal of Plant Science 2020, 11, 653-671.

9. Akter, J.; Jannat, R.; Hossain, M.M.; Ahmed, J.U.; Rubayet, T.M. Chitosan for Plant Growth Promotion and Disease Suppression against Anthracnose in Chilli. International Journal of Environment, Agriculture and Biotechnology 2018, 3, 806–817.

10. Malerba, M.; Cerana, R. Recent Applications of Chitin- and Chitosan-Based Polymers in Plants. Polymers 2019, 11. https://doi.org/10.3390/polym11050839

11. Kulcheski, F.R.; Côrrea, R.; Gomes, I.A.; de Lima, J.C.; Margis, R. NPK macronutrients and microRNA homeostasis. Frontiers in Plant Science 2015, 6. https://doi.org/10.3389/fpls.2015.00451

12. Oluma, H.O.A.; Onekutu, A.; Onyezili, F.N. Reactions of plantain and banana cultivars to black sigatoka leaf spot disease in three farming systems in the Nigerian guinea savanna. Journal of Plant Diseases and Protection 2004, 111, 158–164.

13. Purbajanti, E.D.; Slamet, W.; Fuskhah, E.; Rosyida. Effects of organic and inorganic fertilizers on growth, activity of nitrate reductase and chlorophyll contents of peanuts (Arachis hypogaea L.). IOP Conference Series Earth and Environmental Science 2019, 250. https://doi.org/10.1088/1755-1315/250/1/012048.

14. Kaho, F.; Yemefack, M.; Feudjio-Teguefouet, P.; Tchantchouang, J.C. Effet combiné des feuilles de tithonya diversifolia et des engrais inorganiques sur les rendements du maïs et les propriétés d’un sol ferralitique au Centre Cameroun. Tropicultura 2011, 29, 39–45.

15. Ngosong, C.; Mfombep, P.M.; Njume, C.A.; Tening, A.S. Comparative advantage of *Mucuna* and *Tithonia* residue mulches for improving tropical soil fertility and tomato productivity. International Journal of Plant Soil Science 2016, 12, 1–13.

16. Bilong, E.G.; Ngome, A.F.; Abossolo-Angue, M.; Birang Madong; Ndaka, B.S.M.; Bilong, P. Effets des biomasses vertes de *Tithonia diversifolia* et des engrais minéraux sur la croissance, le développement et le rendement du manioc (*Manihot esculenta* Crantz) en zone forestière du Cameroun. International Journal of Biological and Chemical Science 2017, 11, 1716–1726.

17. Oyerinde, R.O.; Otusanya, O.O.; Akpor, O.B. Allelopathic effect of tithonya diversifolia on the germination, growth and cholorophyl of maize (*Zea mays* L.). Scientific Research and Essay 2009, 4, 879–888.

18. Cobo, J.G.; Barrios, E.; Kaas, D.C.L.; Thomas, R.J. Nitrogen mineralization and crop uptake from surface-applied leaves of green manure species on a tropical volcanic-ash soil. Biology and fertility of soils 2002, 36, 87–92.

19. Diby, Y.K.S.; Tahiri, Y.A.; Akpesse, A.A.M.; Trabi, C.S.; Kouassi, K.P. Evaluation de l’effet insecticide de l’extrait aqueux de Tithonia diversifolia (Hemsl.) gray (Asteracee) sur les termites en culture du riz (NERICA 1) au centre de la Cote d’Ivoire. Journal of Animal & plant Sciences 2015, 25, 3966–3976.

20. Kerebba, N.; Oyedeji, A.O.; Byamukama, R.; Kuria, S.K.; Oyedeji, O.O. Pesticidal activity of *Tithonia diversifolia* (Hemsl.) A. Gray and *Tephrosia vogelii* (Hook f.); phytochemical isolation and characterization: A review. South African Journal of Botany 2019, 121, 366–376.

21. Yuncai, B.; Hucs, Z.; Schmidhalt, U. Effect of Foliar Fertilization Application on the Growth and Mineral Nutrient content of Maize Seedling under Drought and Salinity. Journal of Botany 2008, 5, 1747–1765.

22. Tanaka, Y.; Matsuoka, M.; Yamanoto, N.; Ohashi, Y.; Kano-Murakami, Y.; Ozeki, Y. “Structure and characterization of a cDNA clone for phenylalanine ammonia-lyase from cut-injured roots of sweet potato”. Plant Physiology 1989, 90, 1403–1407.

23. Sharma, A.; Shahzad, B.; Rehman, A.; Bhardwaj, R.; Landi, M.; Zheng, B. Response of Phenylpropanoid Pathway and the Role of Polyphenols in Plants under Abiotic Stress. Molecules 2019, 24. https://doi.org/10.3390/molecules24132452

24. Muymas, P.; Pichyangkur, R.; Wiriyakitnateekul, W.; Wangsomboondee, T.; Chadchawan S.; Seraypheap, K. Effects of chitin-rich residues on growth and postharvest quality of lettuce. Biological Agriculture and Horticulture 2014, 31, 108–117.

25. Kananont, N.; Pichyangkura, R.; Kositsup, B.; Wiriyakitnateekul, W.; Chadchawan, S. Improving the rice performance by fermented chitin waste. International Journal of Agriculture and Biology 2016, 18, 9–15.

26. Lassoudière, A. Le bananier et sa culture. Editions Quae. 2007, 384p.

27. Onautshu, O.D. Caractérisation des populations de *Mycosphaerella fijiensis* et épidémiologie de la cercosporiose noire du bananier (*Musa* spp.) dans la région de Kisangani-République Démocratique du Congo. Thèse de doctorat ès science. Université Catholique de Louvain. 2013, 309p.

28. Ewané, C.A.; Lassois, L.; Brostaux, Y.; Lepoivre, P.; de Lapeyre de Bellaire L. The Susceptibility of Bananas to Crown Rot Disease Is Influenced by Geographic and Temporal Effects. Canadian Journal of Plant Pathology 2012, 35, 27–36.

29. Pirovani, P.C.; Heliana, A.S.C.; Regina, C.R.; Dayane, S.G.; Fatima, C.A.; Fabienne, M. Protein extraction for proteome analysis from cacao leaves and meristems, organs infected by Moniliophthora perniciosa, the causal agent for the witches broom diseases. Electrophoresis journal 2008, 29, 2391–2401.

30. El Hadrami, A. Caractérisation de la résistance partielle des bananiers à la maladie des raies noires et évaluation de la variabilité de l’agressivité de l’agent causal *Mycosphaerella fijiensis*. Thèse présentée en vue de l’obtention d’un Doctorat d’Etat, Gembloux. 2000, 165p.

31. Addinsoft XLSTAT Statistical and data analysis solution. New York, USA. 2020. https://www.xlstat.com.

